# Structures of the human sodium-citrate cotransporter NaCT with and without substrates

**DOI:** 10.64898/2026.07.08.737274

**Authors:** David B. Sauer, Jinmei Song, Jennifer J. Marden, Bing Wang, Kate Sowerby, Joseph C. Sudar, William J. Rice, Da-Neng Wang

## Abstract

The human sodium-citrate cotransporter NaCT imports various tri- and dicarboxylates into the cell as TCA cycle intermediates. This substrate uptake process is driven by an inward sodium gradient. The protein is a member of the Divalent Anion-Sodium Symporter (DASS) family. Whereas extensive biochemical and structural studies have been carried out for NaCT, how the substrate binding and translocation is coupled to the sodium gradient remains unclear. Here using single particle cryo-electron microscopy, we determined the structures of the human NaCT protein in three states: sodium-free, in the presence of sodium, and sodium- and substrate-bound. These structures suggest a simultaneous binding mechanism for sodium-substrate coupling, distinct from the sequential binding, conformational selection mechanism previously observed for the bacterial DASS protein VcINDY.

## 1. Introduction

Citrate plays several important roles in the cell as both an energy source and a major metabolite in the TCA cycle. Not only is the metabolite a major precursor for *de novo* fatty acid biosynthesis ^1^, its concentration in the cytosol upregulates this synthesis pathway by directly binding to the enzyme at the rate-limiting step of the pathway ^2,3^. In the nervous system, where the citrate levels can be as high as 0.4 mM in the cerebrospinal fluid ^4^, the imported tricarboxylate acts as a precursor for neurotransmitter synthesis and can regulate neuronal excitability through the NMDA glutamate receptors ^5–7^. Disrupting citrate catabolism and intracellular citrate accumulation can lead to stress responses and impairs cell fitness ^8^. Therefore, it is critical that the cytosolic levels of citrate are carefully controlled.

The cell has two major sources for obtaining cytosolic citrate, export from the mitochondria and import from the extracellular *milieu* across the plasma membrane. The plasma membrane transporter NaCT (SLC13A5) couples the import of citrate and other di- and tricarboxylates from the extracellular environment to the co-transport of four sodium ions ^9,10^. Various disease phenotypes have been associated with NaCT mutants, including dental hypoplasia and early onset epilepsy ^11–14^. The disease-associated NaCT mutations either reduce the protein’s folding and stability, directly abolish substrate binding, or hinder conformational changes required for translocating the substrate across the membrane ^13,15^. Multiple inhibitors have been developed as starting points or tool molecules for the therapeutic targeting NaCT to treat obesity, kidney diseases and epilepsy ^16–19^, including the substrate mimicking PF2 ^20,21^.

NaCT is a member of the solute carrier 13 gene family (SLC13) of anion transporters ^22–24^. Other members include the sodium-dicarboxylate cotransporters NaDC1 and NaDC3. The mammalian SLC13 proteins are part of the evolutionarily conserved Divalent Anion-Sodium Symporter (DASS) family, with the best characterized member, VcINDY ^25–28^, serving as a prototype for the sodium-coupled anion transport mechanism of the entire family ^29–32^. Over the past few years, significant progress has been made in the mechanistic understanding of the SLC13/DASS proteins, particularly through structure determination ^28^. DASS transporters form homodimers, where each protomer is composed of a scaffold domain and a mobile transport domain, with the substrate and sodium sites located at the middle of the transport domain. The substrate binding site is sandwiched between two sodium sites, Na1 and Na2. Such proximity of the anionic substrate with the sodium ions allows for charge compensation, helping to reduce the energy barrier for substrate translocation across the membrane ^33–35^. For NaDC1 and NaDC3, structures of both the inward-facing (C_i_) and outward-facing (C_o_) conformations have been observed ^30,31^, whereas the NaCT structure has only been determined in the inward-facing conformation and Na^+^-substrate or inhibitor bound states ^29,31^. Such structural information, along with cross-linking experiments, molecular dynamics simulations and single-molecule Förster resonance energy transfer measurements ^27,36,37^ have demonstrated that these SLC13 proteins and their bacterial homologs operate via an elevator mechanism for substrate transport ^38,39^.

Despite such progress, one important mechanistic question for NaCT remains unaddressed, namely: How is substrate transport coupled to sodium binding? The coupling mechanism for substrate and co-substrate in most secondary membrane transporters is poorly understood ^40,41^. Early whole cell transport kinetics and patch-clamp measurements indicate that rabbit NaDC1 employs a sequential binding scheme, with sodium binding first, followed by the dicarboxylate substrate ^42,43^. In this mechanism, it is proposed that the substrate release process mirrors the binding process, wherein the dicarboxylate is released first, followed by the sodium ions.

Transport activity and substrate binding measurements showed that such a sequential binding scheme applies to bacterial DASS transporters as well ^44–46^. Notably, the only DASS protein for which the sodium-substrate coupling mechanism has been extensively characterized is the dicarboxylate transporter from *Vibrio cholerae*, VcINDY. Cryo-EM structure determination of the transporter in different states along with site-directed labelling assays showed that sequential binding is driven by a sodium-induced conformational selection mechanism ^27,35^. In this mechanism, the binding of sodium ions at the Na1 and Na2 sites selects from an ensemble of loops and rigidifies them to a single configuration that can subsequently coordinate the dicarboxylate substrate in VcINDY ^35^.

In addition to citrate, NaCT also transports various dicarboxylates such as succinate and α-ketoglutarate. The transporter binds two sodium ions at the canonical Na1 and Na2 sites of DASS transporters, proximal to the citrate site ^29^. However, it is unknown whether NaCT also follows the same sequential binding, conformational selection mechanism for sodium-substrate coupling. To address this question, we have determined the cryo-EM reconstructions of human NaCT in the absence of sodium, in the presence of sodium, and in sodium and a substrate-mimicking inhibitor. These structures provide high-resolution detail of NaCT’s structure in *apo*, sodium-saturated, and sodium- and dicarboxylate-bound states, and provide an understanding of the protein’s sodium and substrate coupling mechanism.

## 2. Results

Cryo-EM can yield structures of a transporter protein in various substrate-bound states ^31,47^. Applying VcINDY’s conformational selection mechanism for sequential binding ^35^ to NaCT predicts that (1) in the absence of sodium, the human protein *apo* structure is flexible at the Na1 and Na2 sites; (2) binding of sodium stabilizes a discrete, rigid structure that can coordinate the di-/tricarboxylate substrate (S). (3) As a consequence of this mechanism at the Na1/Na2/S sites, the NaCT-Na^+^ structure will differ from the *apo* state but will resemble the NaCT-Na^+^-S state. To examine these predictions, we need the structures of NaCT-Na^+^-S, NaCT-Na^+^, and NaCT-*apo*.

### 2.1. NaCT-Na^+^-PF2 structure

In our previous studies, structures of NaCT-Na^+^-S have been determined ^29,31^ in the presence of 100 mM NaCl, together with one of the four substrates, citrate, succinate, PF2 (PF-06649298) and PF4a (PF-067611281), with the latter two being substrates that enter the cell via NaCT, bind from the cytosol and become competitive inhibitors ^20,21^. Under all conditions tested, the transporter dimer adopts a symmetric, inward-facing C_i_-C_i_ conformation. However, precise analysis of the protein structure and ion binding in these structures is challenged due to the weak affinity of DASS transporters for sodium (K_0.5_ ∼70 mM) ^48–51^, which would result in low occupancy for sodium ions at 100 mM within the protein. The density for sodium is likely diminished further by the 20 mM LiCl included in the cryo-EM sample preparation, where this lighter ion thermostabilizes the protein ^29^ but gives a weaker signal in the Coulombic potential map ^52^.

To resolve this ambiguity in sodium visibility in NaCT maps, we purified NaCT in the presence of both 300 mM NaCl and 100 μM PF2, but in the absence of LiCl. This sodium concentration of 300 mM was chosen to balance the desire for high occupancy with the need to avoid any salting-out effects on the protein in solution and to minimize background noise in the electron micrographs ^27,53,54^. PF2 was chosen over citrate in the structure determination for higher affinity and occupancy ^29^. The detergent-purified protein was exchanged into the amphipol polymer and cryo-EM grids were frozen. The NaCT particles on the grids displayed preferred orientation problems as was observed with previous samples prepared in 100 mM NaCl, which we once again overcame by collecting electron micrographs at 0 and 40° tilts ^29,55^. Processing the merged datasets allowed us to solve the NaCT structure at an overall resolution of 2.53 Å (Figs. 1A & B, Fig. S1). The protein adopted a C_i_-C_i_ dimer structure (Fig. 1C), with densities for PF2, Na1 and Na2 confirming that NaCT was captured in the C_i_-Na^+^-PF2 state (Fig. 1D-F). The overall structure is similar to that previously solved at 3.12 Å resolution in 100 mM NaCl, 20 mM LiCl and 100 μM PF2 ^29^ (R.M.S.D. = 0.40 Å). Nevertheless, the higher experimental resolution and clearer densities at the PF2, Na1, and Na2 binding sites enabled us to build a more accurate structural model for the protein and its bound cations and substrate (Table 1). This PF2-bound structure is very similar to the one previously-determined in citrate ^29^ (R.M.S.D. = 0.79 Å), and we thus conclude it represents the substrate-bound NaCT-Na^+^-S state.

**Figure 1.**
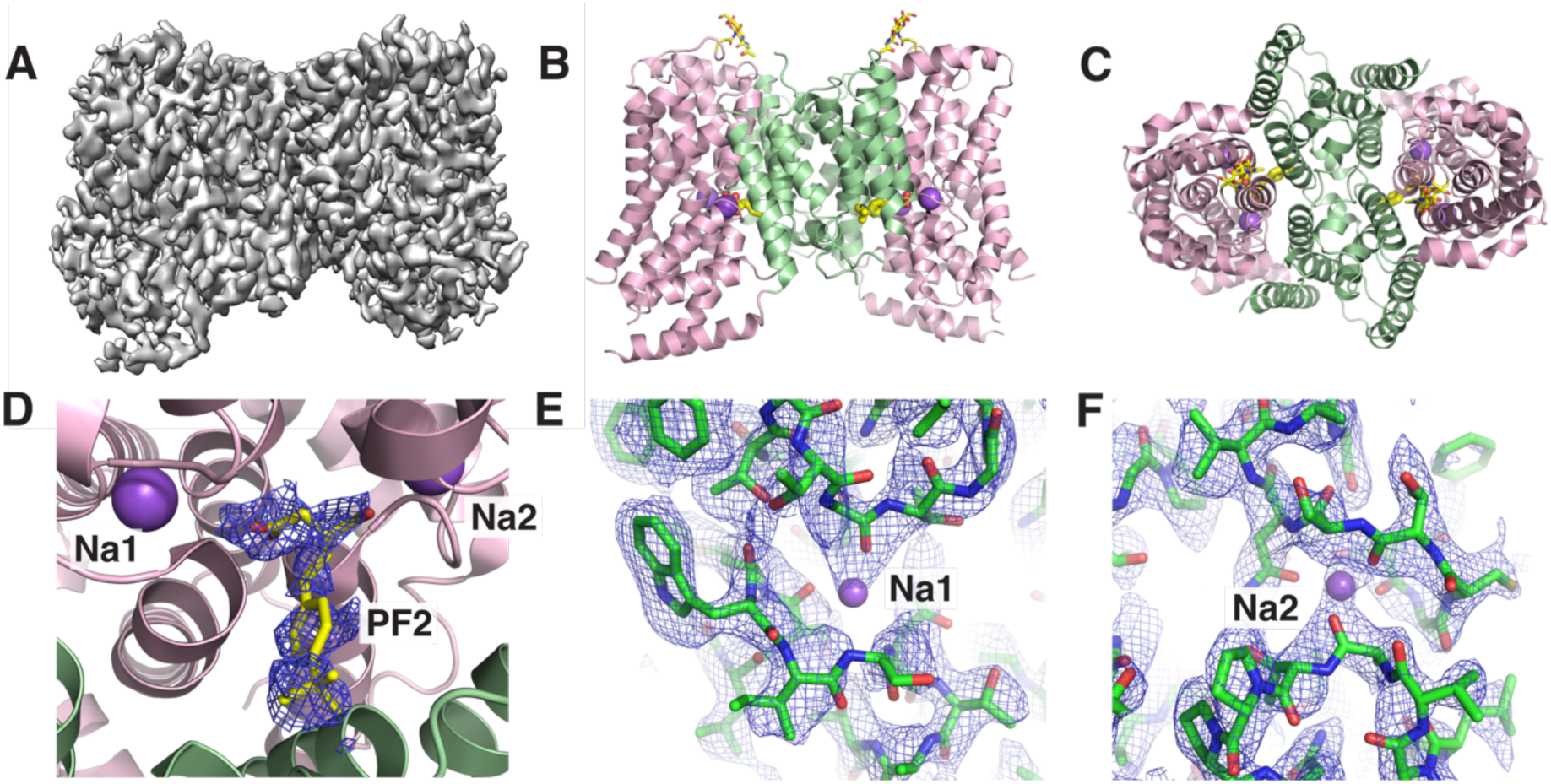
High-resolution structure of sodium and PF2-bound NaCT. (A) Columbic potential map of NaCT determined to 2.53 Å resolution. Atomic model of NaCT viewed from (B) within the membrane and (C) outside the cell. (D) The PF2-binding site structure with Columbic potential map is shown in mesh. The scaffold and transport domains are colored light green and light pink, respectively. The PF2 substrate-mimicking inhibitor is shown in yellow. The (E) Na1 and (F) Na2 binding sites with the cryo-EM map shown as mesh and contoured at 10 σ.

**Table 1.** Cryo-EM data collection, image processing and structure determination.

|  | **NaCT-Na<sup>+</sup>-PF2** | **NaCT-Na<sup>+</sup>** | **NaCT-*apo*** |
| --- | --- | --- | --- |
| **EMDB Code** | EMD-58071 | EMD-58072 | EMD-58073 |
| **PDB Code** | 30UT | 30UU | 30UV |
| **Microscope** | NYSBC Krios | NYU Krios | NYU Krios |
| **Detector** | K3 | K3 | K3 |
| **Voltage (kV)** | 300 | 300 | 300 |
| **Magnification** | 105,000 | 105000 | 105000 |
| **Collection mode** | Superresolution | Superresolution | Superresolution |
| **Electron exposure (e/Å<sup>2</sup>)** | 63.34 | 53.61 | 57.95 |
| **Number of frames** | 50 | 48 | 48 |
| **Pixel size (Å)** | 0.826 | 0.825 | 0.852 |
| **Defocus range (µm)** | -4.0 to -6.9 | -0.5 to -2.5 | -1 to -2 |
| **Number of movies** |  |  |  |
| 0° tilt | 1585 | 2353 | 2237 |
| 40° tilt | 2464 | 3192 | 5473 |
| **Initial Number of particles** | 2,620,654 | 2,335,793 | 3,170,073 |
| **Symmetry** | C2 | C2 | C2 |
| **Number of particles used for final 3D refinement** | 976,961 | 951,970 | 1,503,602 |
| **Map resolution (Å ; FSC threshold = 0.143)** | 2.53 | 2.51 | 2.49 |
| **Model resolution (Å ; FSC threshold = 0.5)** | 2.8 | 3.0 | 2.7 |
| **Non-hydrogen atoms** | 7388 | 7342 | 7342 |
| Protein residues | 936 | 936 | 936 |
| Ligands | 8 | 2 | 2 |
| **R.M.S.D** |  |  |  |
| Bond lengths (Å) | 0.003 | 0.003 | 0.003 |
| Bond angles (°) | 1.056 | 1.075 | 1.053 |
| **Validation** |  |  |  |
| Molprobity score | 2.28 | 1.91 | 1.91 |
| Clash score | 7.56 | 4.99 | 4.19 |
| Rotamer outliers (%) | 3.74 | 1.75 | 2.37 |
| **Ramachandran plot** |  |  |  |
| Favoured (%) | 93.45 | 92.58 | 93.56 |
| Allowed (%) | 6.55 | 7.31 | 6.33 |
| Disallowed (%) | 0.00 | 0.11 | 0.11 |

### 2.2. Structures of sodium-bound and *apo* NaCT

To identify structural changes in NaCT through its reaction cycle, we next determined the structures of NaCT in the absence of substrate, or in the absence of both substrate and sodium, to capture the NaCT-Na^+^ and NaCT-*apo* states, respectively. The NaCT-Na^+^ sample was prepared in the presence of 300 mM NaCl, whereas the *apo* sample was purified in 100 mM choline chloride (ChCl). Choline has been shown to be an organic cation that can maintain the integrity and stability of Na^+^ channels and Na^+^-driven transporters during purification ^35,56,57^. As with the previous sample in NaCl and PF2, NaCT particles on both the 300 mM NaCl and 100 mM ChCl grids exhibited preferred orientations. Nevertheless, using data collection from tilted grids ^29,55^, we were able to obtain 3D maps and protein models of the protein at overall isotropic resolutions of 2.51 Å and 2.49 Å (Table 1, Figs. S2&S3), for sodium-saturated and sodium-free samples, respectively (Figs. 2A&B, 2E&F). No density was seen at the citrate, Na1 or Na2 sites for NaCT in 100 mM ChCl (Figs. 2G&H), confirming that we have captured the protein in an *apo* state.

**Figure 2.**
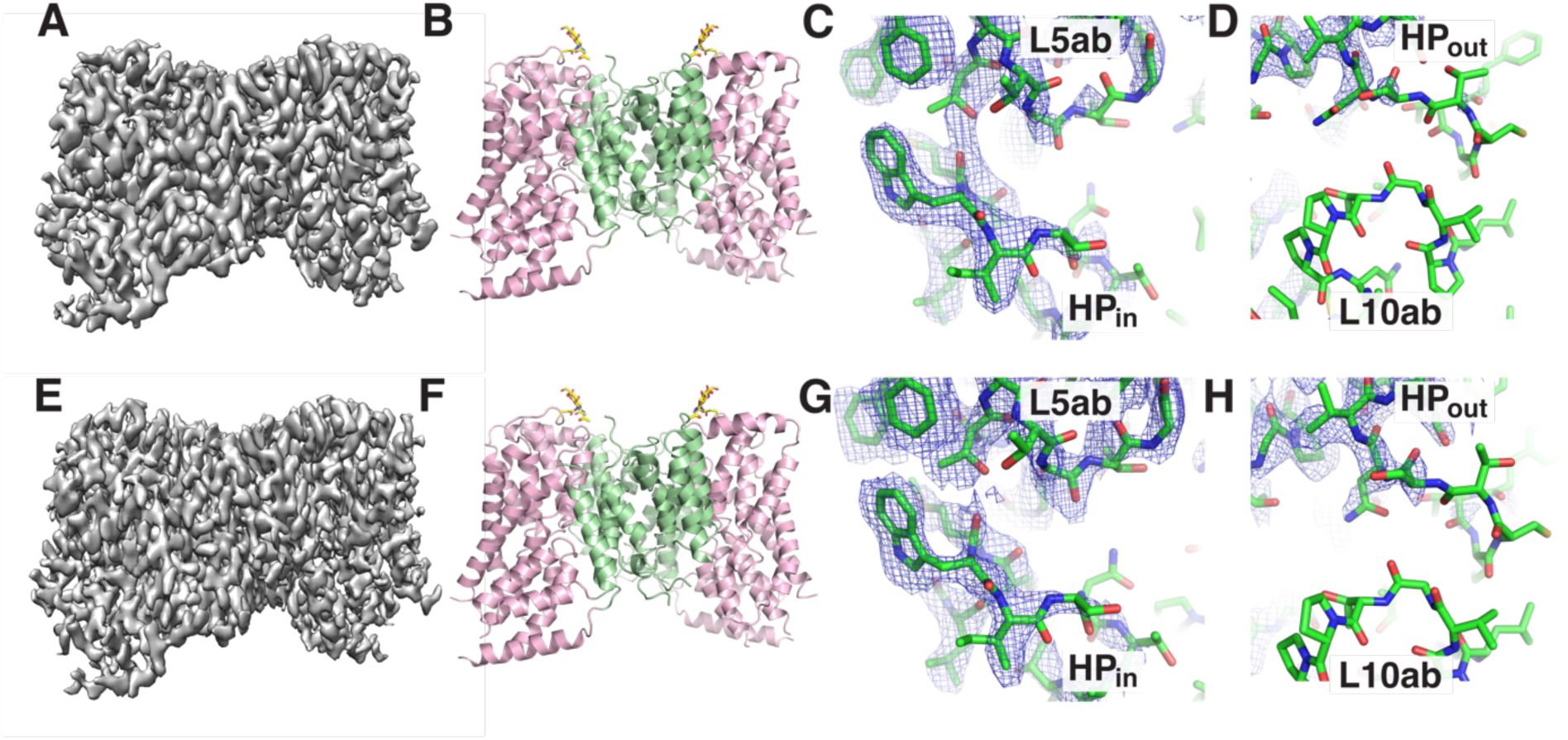
Structure of NaCT’s C_i_-Na^+^ and C_i_-*apo* states. (A) The Columbic potential map and (B) atomic model of NaCT-Na^+^. (C) The Na1 and (D) Na2 binding sites for the NaCT-Na^+^ state with the cryo-EM map shown as mesh and contoured at 10 σ. (E) The Columbic potential map and (F) atomic model of NaCT-*apo*. (G) The Na1 and (H) Na2 binding sites for the NaCT-*apo* state with the cryo-EM map shown as mesh and contoured at 10 σ.

Remarkably, we did not observe any density at the Na1 and Na2 sites for NaCT in 300mM NaCl (Figs. 2C&D), and the protein density near the Na2 site was of poor quality in both conditions. The absence of cation densities at the Na1and Na2 sites in the sodium-only condition and the increased protein mobility near Na2, are both marked differences from VcINDY in equivalent conditions ^35^ and suggest a different cation-substrate coupling mechanism for NaCT.

Structures of sodium-saturated and *apo* NaCT both adopted a symmetric, inward-facing conformation, termed C_i_-Na^+^ and C_i_-*apo*, respectively. Notably, while the C_i_-Na^+^ and C_i_-*apo* structures were essentially identical (R.M.S.D. = 0.48 Å) (Fig. 3A), there were marked changes between C_i_-Na^+^ and C_i_-Na^+^-PF2 structures (R.M.S.D. = 1.1 Å). This is most pronounced on the horizontal helices H4c and H6b (Fig. 3A), and apparent movements of HP_in_, and TM10b on the cytoplasmic face of the transport domain (Fig. 3B), necessitating a detailed exploration of how PF2 release triggers these subtle structural movements.

**Figure 3.**
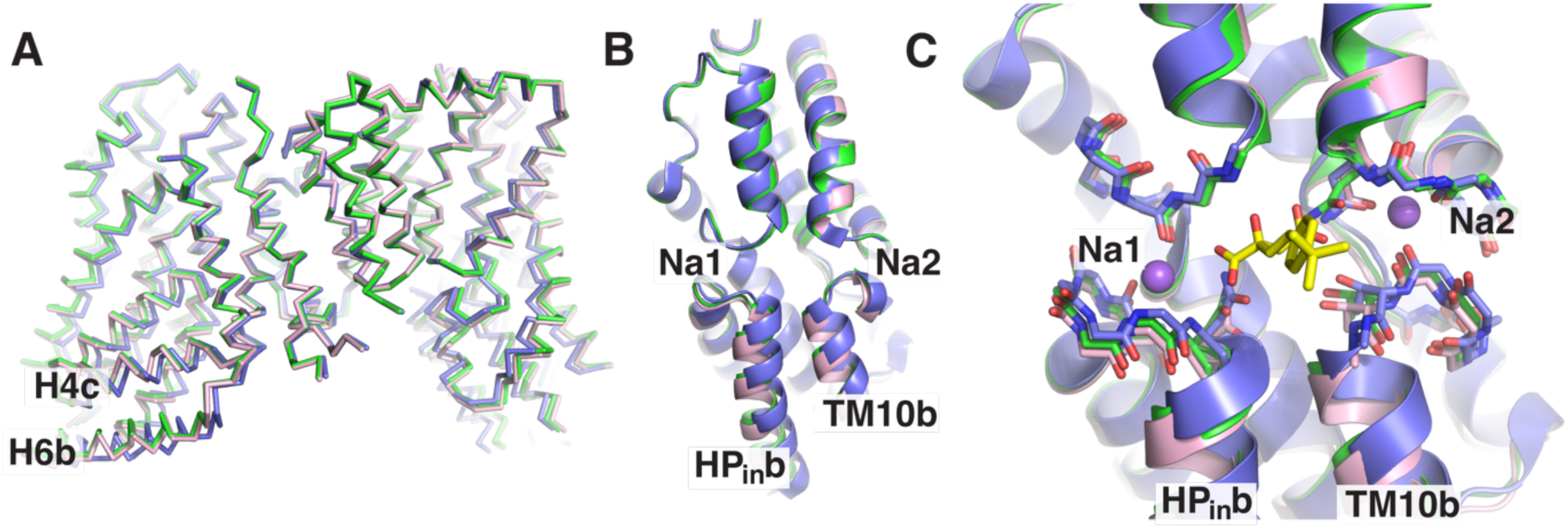
Carboxylate release triggers conformational changes in the NaCT transport domain. A) Movements in the NaCT transport domain and arm helices H4c and H6b upon PF2 release. C_i_-Na^+^-PF2, C_i_-Na^+^, and C_i_-*apo* states are colored in blue, green, and pink, respectively. PF2 release triggers movements in (B) HP_in_ and TM10b due to changes in the (C) sodium and carboxylate binding site.

Examining the substrate and sodium binding sites, it is immediately apparent that the structural changes observed between C_i_-Na^+^ and C_i_-Na^+^-PF2 is triggered by NaCT-substrate/inhibitor interactions (Fig. 3C). In particular, PF2 makes hydrogen bonds with Asn141, Thr142, Thr227, Gly228, Asn465 and Thr508, bridging the core of the transport domain and HP_in_ and TM10b. Consequently, release of carboxylate substrate relieves this structural restraint to trigger the structural movements of HP_in_ and TM10b that are subsequently propagated to the interfacial helices through the hinge loop connecting HP_in_a to H6a. Notably, such differences between the C_i_-Na^+^ and C_i_-Na^+^-PF2 structures are inconsistent with the predictions made by the sequential binding, conformation selection coupling mechanism ^35^.

### 2.3. Binding site flexibility upon substrate release

While the cryo-EM maps were sufficient to build atomic models for the entire NaCT transporter in all states, markedly weaker density was observed near Na2 in the C_i_-Na^+^ and C_i_-*apo* states (Fig. 2). Furthermore, we recognized that, while traditional model refinement provides a single protein model for each experimental condition, the varying strength of the experimental density might reflect differences in mobility across the protein structure. We therefore next examined changes in NaCT flexibility between the three conditions by directly comparing the cryo-EM densities, along with the convergence and atomic mobility in the refined models.

Examining the equivalently scaled Coulombic potential maps, we immediately recognized that the density for TM10b of NaCT was poorly resolved for the C_i_-Na^+^ and C_i_-*apo* states, relative to C_i_-Na^+^-PF2 (Figs. 4A-C), even though the densities for the rest of the protein were well defined. This suggested greater structural flexibility of TM10b near Na2 in the absence of substrate; as alternative mechanisms for the observed map differences, such as radiation damage or poor particle alignment, would not result in region-specific effects. This similarity between the C_i_-Na^+^ and C_i_-*apo* structures, and differences between the C_i_-Na^+^ and C_i_-Na^+^-PF2 structures, are again in conflict with the predictions of sequential binding via a conformational selection mechanism ^35^.

**Figure 4.**
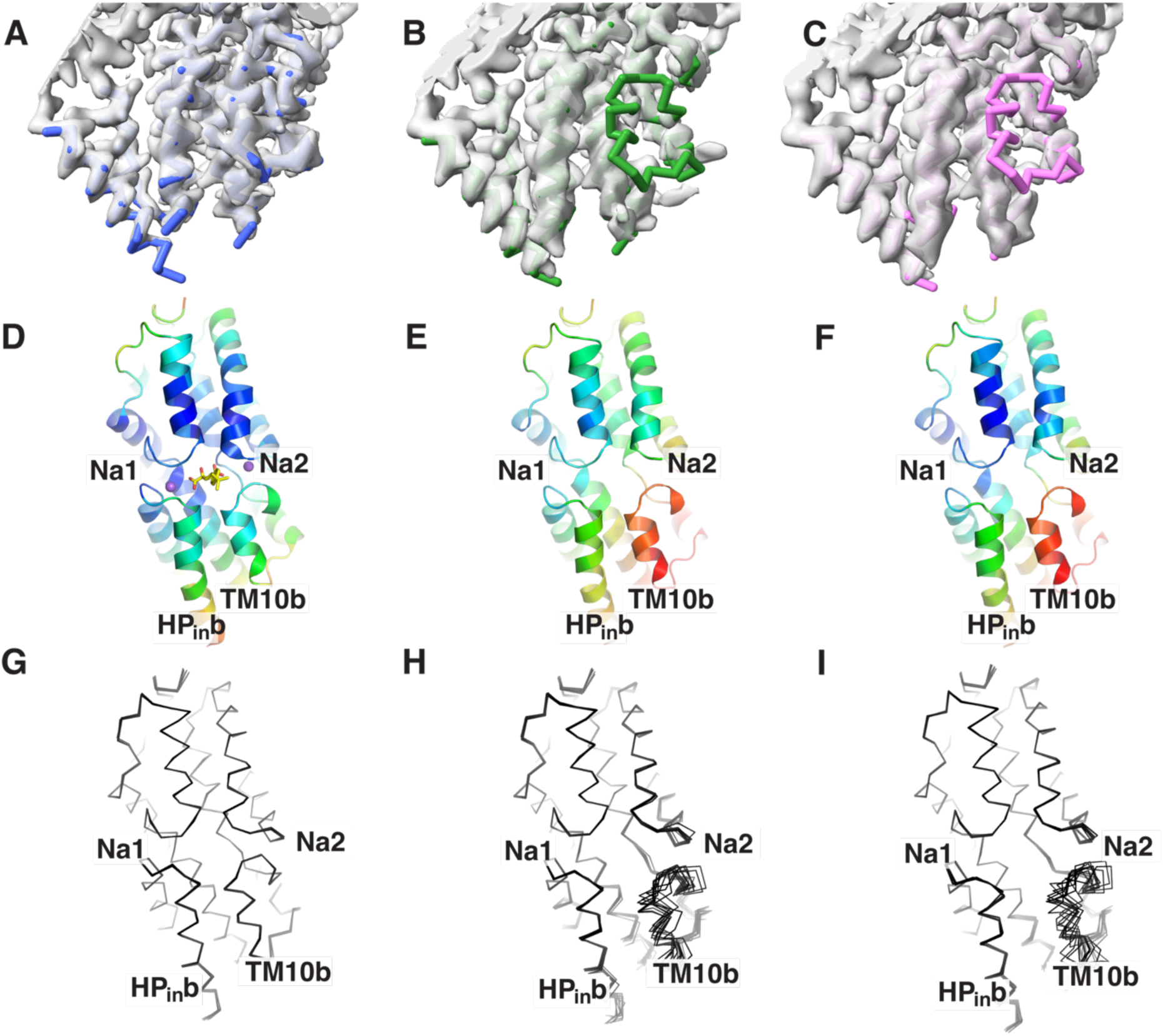
Transport domain flexibility changes between NaCT’s C_i_-Na^+^-PF2, C_i_-Na^+^, and C_i_-*apo* states. Equivalently scaled unsharpened cryo-EM maps of the (A) C_i_-Na^+^-PF2, (B) C_i_-Na^+^, and (C) C_i_-*apo* states, with refined atomic models shown in blue, green, and pink, respectively. Structures of the (D) C_i_-Na^+^-PF2, (E) C_i_-Na^+^, and (F) C_i_-*apo* states, colored by the normalized model B-factors. NMR-style analysis of the (G) C_i_-Na^+^-PF2, (H) C_i_-Na^+^, and (I) C_i_-*apo* states from 10 independent refinements.

Having identified this region of state-dependent flexibility in NaCT, we next sought to capture these structural dynamics in the states’ corresponding atomic models. Although the refined models of NaCT across states were generally similar, atomic B-factors were elevated near the Na2 site of the C_i_-Na^+^ and C_i_-*apo* models (Figs. 4D-F). However, we suspected a single atomic model might not adequately capture the structural ensemble of NaCT in each experimental map and the corresponding state. Therefore, we conducted an NMR-style analysis using multiple independent rounds of simulated annealing and model refinement ^35^. In this method, the rigid parts of the protein will converge to the same coordinates, while after each run mobile regions will be at distinct atomic positions that fits equally into the map. As expected, the models converged across most of the well-resolved regions of NaCT protein and its transport domain (Figs. 4G-I). However, significant structural heterogeneity was observed near the Na2 site and in TM10b in the absence of PF2.

## 3. Discussion

VcINDY’s conformational selection model provided the first mechanistic model for energy-coupling by a DASS family member ^35^. However, significant sequence and functional differences between the prokaryotic VcINDY and human NaCT suggested the human protein might not use an identical mechanism to couple cation and carboxylate binding. Here, we determined structures of NaCT in its C_i_-Na^+^-PF2, C_i_-Na^+^ and C_i_-*apo* states to understand this eukaryotic protein’s coordination of sodium and substrate-mimicking inhibitor, PF2.

In the absence of sodium, both NaCT and VcINDY show greater flexibility in the Na2 site and TM10b helix than the Na1 site. Remarkably, the substrate/sodium-induced structural changes of NaCT differ significantly from those observed in its bacterial homolog VcINDY. First, NaCT is rigid at the Na1 site in the absence of sodium, whereas the structure of VcINDY is highly flexible in the absence of a cation. Second, the presence of sodium was sufficient to fix the loop positions at the Na2 site in VcINDY for dicarboxylate binding, whereas this site and TM10b in NaCT become rigid only in the presence of both sodium and a carboxylate substrate. In summary, for VcINDY the binding site structures are similar between C_i_-Na^+^-S and C_i_-Na^+^ and different between C_i_-Na^+^ and C_i_-*apo*, indicating that sodium ions alone can select a conformation for dicarboxylate binding. In contrast, NaCT’s C_i_-Na^+^ and C_i_-*apo* share similar binding site structures, which are different from those of C_i_-Na^+^-S. This shows that sodium alone is insufficient to select a configuration in NaCT for substrate binding. These contradictions demonstrate that NaCT couples sodium and substrate binding by a mechanism different from VcINDY’s sodium-dependent conformational selection.

The origin of NaCT and VcINDY’s distinct sodium and substrate coupling mechanisms are unclear due to differences in sequence and function between the proteins. The proteins are distant homologs, with ∼30% pairwise amino acid sequence identity. In addition, NaCT binds 4 sodium ions ^10,50^, versus VcINDY’s 3 co-transported cations ^51^, which may be important to the coupling mechanism. Furthermore, while the sodium binding sites Na1 and Na2 have been identified and are structurally homologous between both proteins, the remaining cation sites remain unknown. Frustratingly, our high-resolution structures of NaCT in saturating sodium conditions do not give any indication of the locations of these additional ions.

Finally, the sequential binding and co-transport of sodium and carboxylates by DASS proteins are typically studied through their import reactions. Under these conditions, sodium and substrate binding occur when the protein is in an outward-facing state and in the presence of a 10-fold transmembrane sodium gradient. In contrast, our results describe the cytoplasmic release of carboxylate and sodium. While the pseudosymmetry of the DASS sequence and structure ^25,36^, and the transporters’ reversibility ^22^, suggest substrate and cation interactions in the inward-facing and outward-facing reactions are generally equivalent, this might not hold true for every detail of the proteins’ reaction cycle or for every protein in the family. Fully exploring this will require capturing a complete reaction cycle, including outward-facing states, for both DASS prototype VcINDY and the human NaCT.

Tentatively, the structural differences between NaCT’s C_i_-Na^+^-PF2, C_i_-Na^+^, and C_i_-*apo* states suggest one possible mechanism for substrate and sodium coupling. NaCT’s carboxylate coordinating residues are directly linked to the Na1 and Na2 binding sites. Consequently, the human protein’s cation binding may be substrate dependent, with the occluded Na1 and Na2 sites in our C_i_-Na^+^-PF2 structure suggesting the sodium and carboxylate binding is simultaneous. This effectively maintains the transporter in an *apo* state unless both sodium and carboxylate substrate are present, and thereby prevents non-productive leakage where only the cation is transported across the membrane. Such a coupling mechanism satisfies the charge compensation requirement for translocating highly-charged substrates across the membrane.

## Resource availability

### Lead contact

Further information and requests for resources and reagents should be directed to and will be fulfilled by the lead contact, Da-Neng Wang.

### Materials availability

All unique reagents generated in this study are available from the lead contact with a completed Materials Transfer Agreement.

### Data and code availability

- Cryo-EM maps and models have been deposited to Protein DataBank, and are publicly accessible. NaCT-Na^+^-PF2 structure map and model have the following accession codes: 30UT (PDB), EMD-58071 (EMDB). NaCT-Na^+^ structure map and model have the following accession codes: 30UU (PDB), EMD-58072 (EMDB). NaCT-*apo* structure map and model have the following accession codes: 30UV (PDB). EMD-58073 (EMDB).
- This paper does not report original code.
- Any additional information required to reanalyze the data reported is available from the lead contact upon request.

## Acknowledgements

This work was financially supported by the NIH (R01NS108151, R01DK135088 and R01GM121994), the G. Harold and Leila Y. Mathers Foundation, and the TESS Research Foundation. D.B.S. was supported by the American Cancer Society Postdoctoral Fellowship (129844-PF-17-135-01-TBE) and the Department of Defense Horizon Award (W81XWH-16-1-0153). The cryo-EM data was collected at the NYU Cryo-Electron Microscopy Core Facility (RRID: SCR_019202), which is partially supported by the Laura and Isaac Perlmutter Cancer Center Support Grant NIH/NCI P30CA016087, and at the Simons Electron Microscopy Center at the New York Structural Biology Center, which is supported by the Simons Foundation (SF349247). We are grateful to E. Eng for technical support. EM data processing used computing resources at the HPC Facility of NYULMC.

## Author contributions

J.S. expressed and purified the protein. J.J.M., K.S., and J.C.S. conducted biochemical studies. D.B.S. froze grids. B.W. and W.J.R. collected and processed the cryo-EM images. D.B.S. built the atomic models. D.B.S and D.N.W. analyzed the structures and wrote the manuscript. All authors participated in the discussion and manuscript editing. D.N.W. supervised the research.

## STAR*Methods

### KEY RESOURCES TABLE

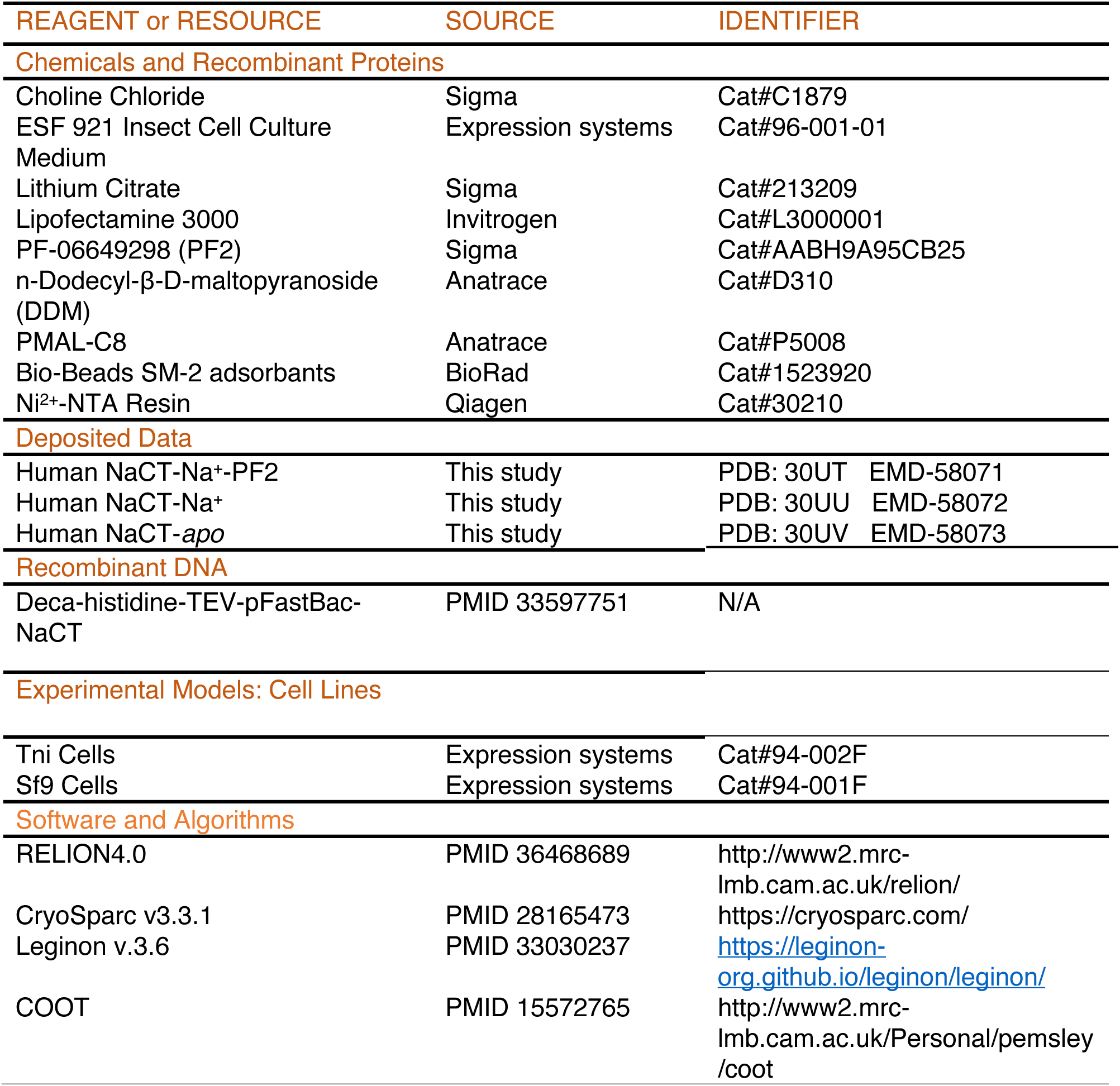
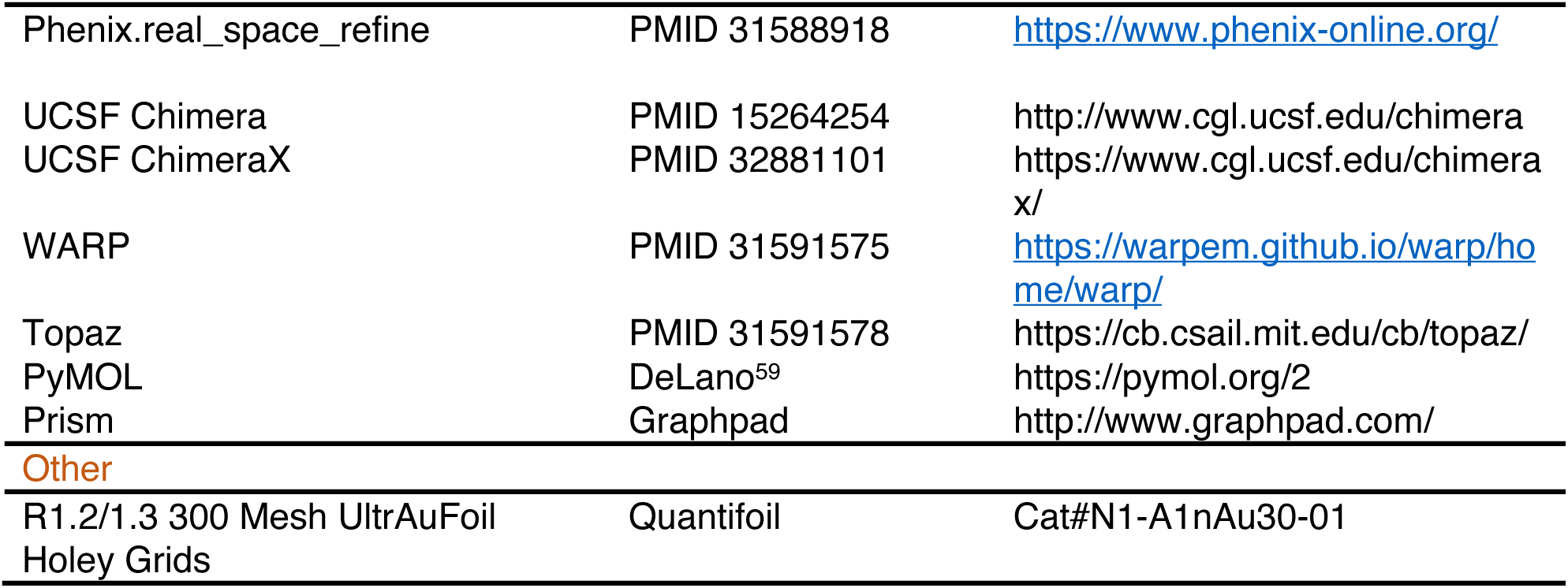

## Method Details

### Overexpression and purification

NaCT was expressed and purified as previous described with minor modifications ^29^. Briefly, T. ni BTI-Tn-5B1-4 cells were transduced with baculovirus bearing the wild-type SLC13A5 gene with an N-terminal decahistidine-TEV tag. Cells were lysed in a buffer of 50 mM Tris pH 8.0, 400 mM NaCl, 10 mM lithium citrate and 10% glycerol. Membranes were collected by ultracentrifugation and solubilized in a buffer of 50 mM Tris pH 8.0, 200 mM NaCl, 10 mM lithium citrate, 10% glycerol and 1.3% DDM. NaCT was subsequently purified on a Ni^2+^-NTA affinity column and exchanged into the amphipol PMAL-C8 (Anatrace). DDM was removed with Bio-Beads SM2 (Bio-Rad). Samples were further purified by size-exclusion chromatography in buffer containing 25 mM Tris pH 7.5, 0.1 mM TCEP and either 300 mM NaCl and 100 μM PF2 (for NaCT-Na^+^-PF2) or 300 mM NaCl (for NaCT-Na^+^) or 100 mM choline chloride (NaCT-*apo*).

### Cryo-EM specimen preparation and image processing

All cryo-EM grids were prepared by applying 3 μl of protein at around 0.75 mg ml^−1^ to a glow-discharged UltrAuFoil R1.2/1.3 300-mesh grid (Quantifoil) and blotted for 2.5–4 s under 100% humidity at 4°C before plunging into liquid ethane using a Mark IV Vitrobot (FEI).

Micrographs of the NaCT–Na^+^-PF2 complex were acquired on a Titan Krios microscope with a K3 direct electron detector and an energy filter slit width of 20 eV. The magnification used was 105,000×, with a physical pixel size of 0.826 Å. Leginon ^58^ was used to for ice thickness targeting, measured as described previously ^59^, resulting in 1,585 and 2,464 micrographs being collected from 0° and 40° tilted specimens, respectively. Each micrograph was dose-fractioned over 50 frames, with an accumulated dose of 63 e^−^ Å^−2^. Frame alignment was performed with Relion 4 ^60^.

Aligned and dose weighted micrographs were imported to Warp ^61^. After particle picking and CTF estimation in WARP, micrographs with an overall resolution worse than 12 Å were excluded from subsequent steps. Initial Warp particle picks were cleaned up in CryoSparc ^62^ with two-dimensional and three-dimensional classification, and then used for repicking using Topaz ^63^ to yield 2,620,564 particles from all micrographs. In CryoSparc, these particles were subjected to multiple rounds of three-dimensional classification to produce a set of 1,000,432 particles, which generated a 3.11 Å reconstruction in nonuniform refinement with C2 symmetry imposed. This map and particle set were then used for Relion ^64^ particle polishing. The particles were re-imported in CryoSparc for global and local CTF optimization and subsequent nonuniform refinement to yield a final 2.53 Å reconstruction with C2 symmetry imposed.

Micrographs of the NaCT-Na^+^ state were acquired on a Titan Krios microscope with a K3 direct electron detector and an energy filter slit width of 20 eV. The magnification used was 105,000×, with a super-resolution pixel size of 0.426 Å. Leginon was used to target holes with 10 to 70 nm of ice, measured as described previously, resulting in 2,353 and 3,192 micrographs being collected from 0° and 40° tilted specimens, respectively. Each micrograph was dose-fractioned over 48 frames, with an accumulated dose of 53.6 e^−^ Å^−2^. Frame alignment was performed with MotionCorr2 ^65^ under control of Appion. Aligned, dose-weighted micrographs were imported to Warp. After particle picking and CTF estimation in Warp, micrographs with an overall resolution worse than 8 Å were excluded from subsequent steps, with the initial set comprising 2,349,354 particles. Warp-picked particle sets from the tilted and untilted micrographs were cleaned using CryoSparc’s two-dimensional and three-dimensional classification, and then combined. These 1,421,195 particles were subjected to multiple rounds of three-dimensional classification to produce a set of 951,970 particles, which generated a 2.63 Å reconstruction in nonuniform refinement with C2 symmetry imposed. This map and particle set were then used for Relion particle polishing. The particles were re-imported in CryoSparc for subsequent nonuniform refinement to yield a final 2.51 Å reconstruction with C2 symmetry imposed.

Micrographs of the NaCT-*apo* state were acquired on a Titan Krios microscope with a K3 direct electron detector and an energy filter slit width of 20 eV. The magnification used was 105,000×, with a super-resolution pixel size of 0.426 Å. Leginon was used to target holes with 15 to 95 nm of ice, measured as described previously, resulting in 2,237 and 5,473 micrographs being collected from 0° and 40° tilted specimens, respectively. Each micrograph was dose-fractioned over 48 frames, with an accumulated dose of 58.0 e^−^ Å^−2^. Frame alignment was performed with MotionCorr2 ^65^ under control of Appion, and aligned. Dose-weighted micrographs were imported to Warp. After particle picking and CTF estimation in Warp, micrographs with an overall resolution worse than 8 Å were excluded from subsequent steps. Initial Warp picks of 1,117,117 particles were cleaned up using CryoSparc’s two-dimensional and three-dimensional classification, and then used for repicking using Topaz to yield 3,170,073 particles from all micrographs. In CryoSparc, these particles were subjected to multiple rounds of three-dimensional classification to produce a set of 1,503,602 particles, which generated a 2.73 Å reconstruction in nonuniform refinement with C2 symmetry imposed. This map and particle set were then used for Relion particle polishing. The particles were re-imported in CryoSparc for subsequent nonuniform refinement to yield a final 2.49 Å reconstruction with C2 symmetry imposed.

### Atomic models were built in Coot ^66^ and refined with Phenix.real_space_refine ^67^

The NMR-style analysis was performed as previously described ^35^, using 5 independent runs of phenix.real_space_refine to refine the models, with ions and ligands removed, against the unsharpened maps with a unique computational seed for each run. Each refinement was performed with simulated annealing, without NCS constraints or secondary structure restraints, and a refinement resolution limit of 2.53 Å for all maps and models. When comparing the local density, unsharpened maps were scaled to equivalent contours using the scaffold domain’s volume as an internal standard after extracting with phenix.map_box ^67^. Figures were made using UCSF Chimera ^68^, ChimeraX ^69^, and PyMOL ^70^.

