## Supplementary material for "Structures of the human sodium-citrate cotransporter NaCT with and without substrates": SI materials

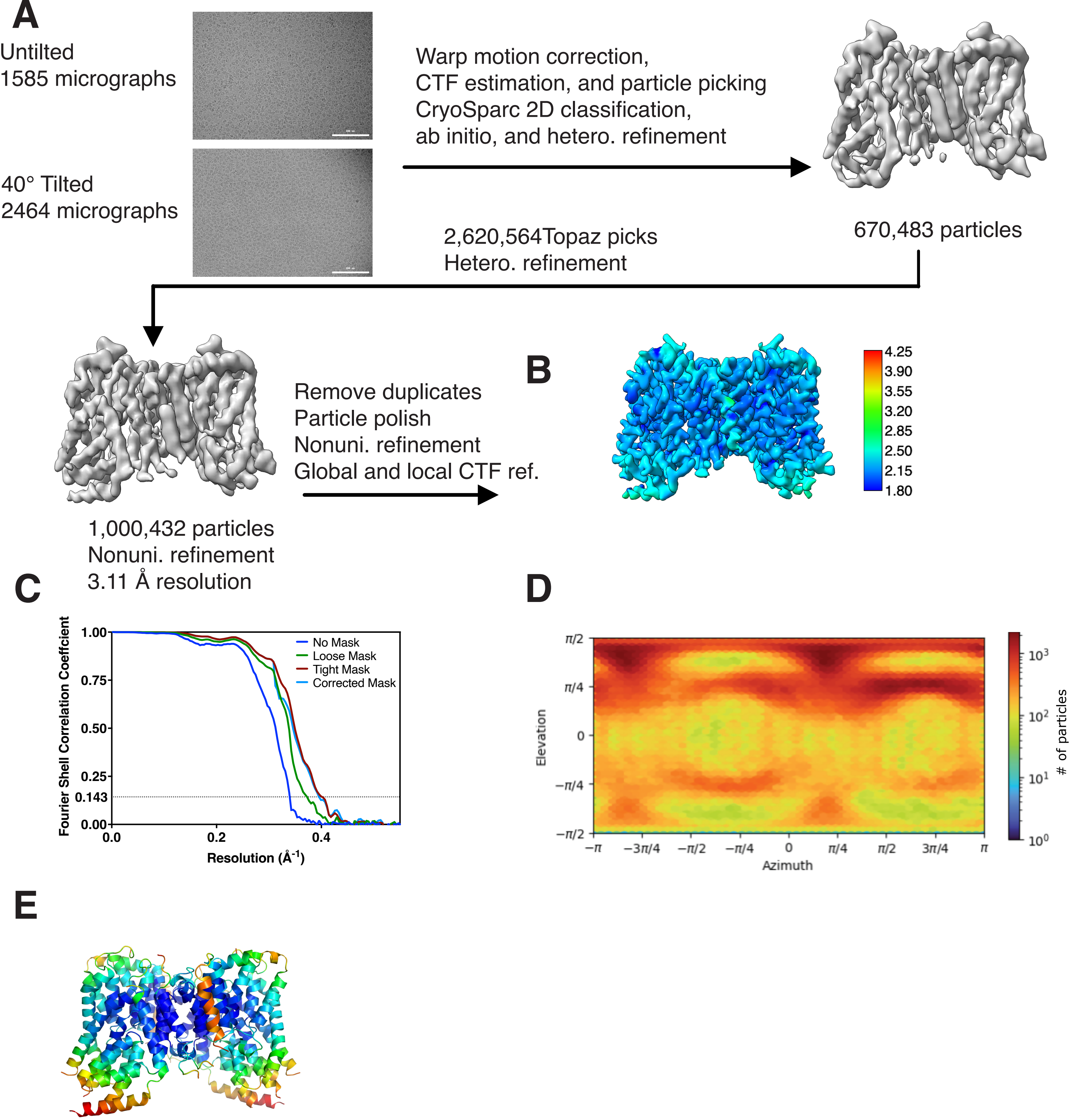


**Supplementary Figure 1. Cryo-EM data processing of the NaCT-Na+-PF2 structure.**
A. Cryo-EM data processing workflow for the NaCT-Na+-PF2 dataset. B. Local resolution map of the NaCT-Na+-PF2 map. C. FSC curve for the NaCT-Na+-PF2 map. D. Angular distribution of particles in the NaCT-Na+-PF2 map. E. Model of NaCT-Na+-PF2, colored by the relative atom B-factors.

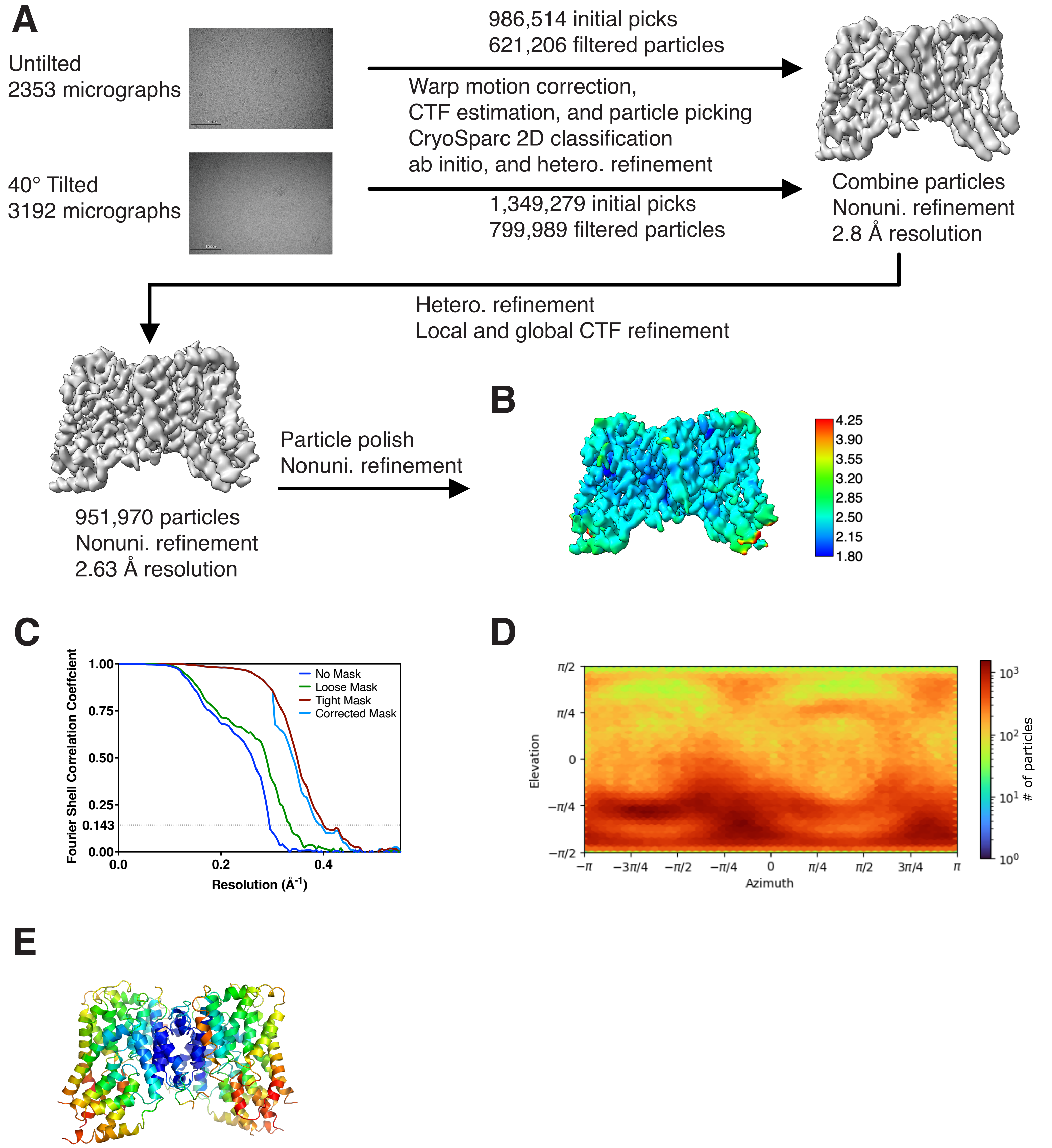


**Supplementary Figure 2. Cryo-EM data processing of the NaCT in NaCl structure.** A. Cryo-EM data processing workflow for the NaCT-Na dataset. B. Local resolution map of NaCT in NaCl. C. FSC curve for the NaCT in NaCl map. D. Angular distribution of particles in the NaCT in NaCl map. E. Model of NaCT NaCl, colored by the relative atom B-factors.

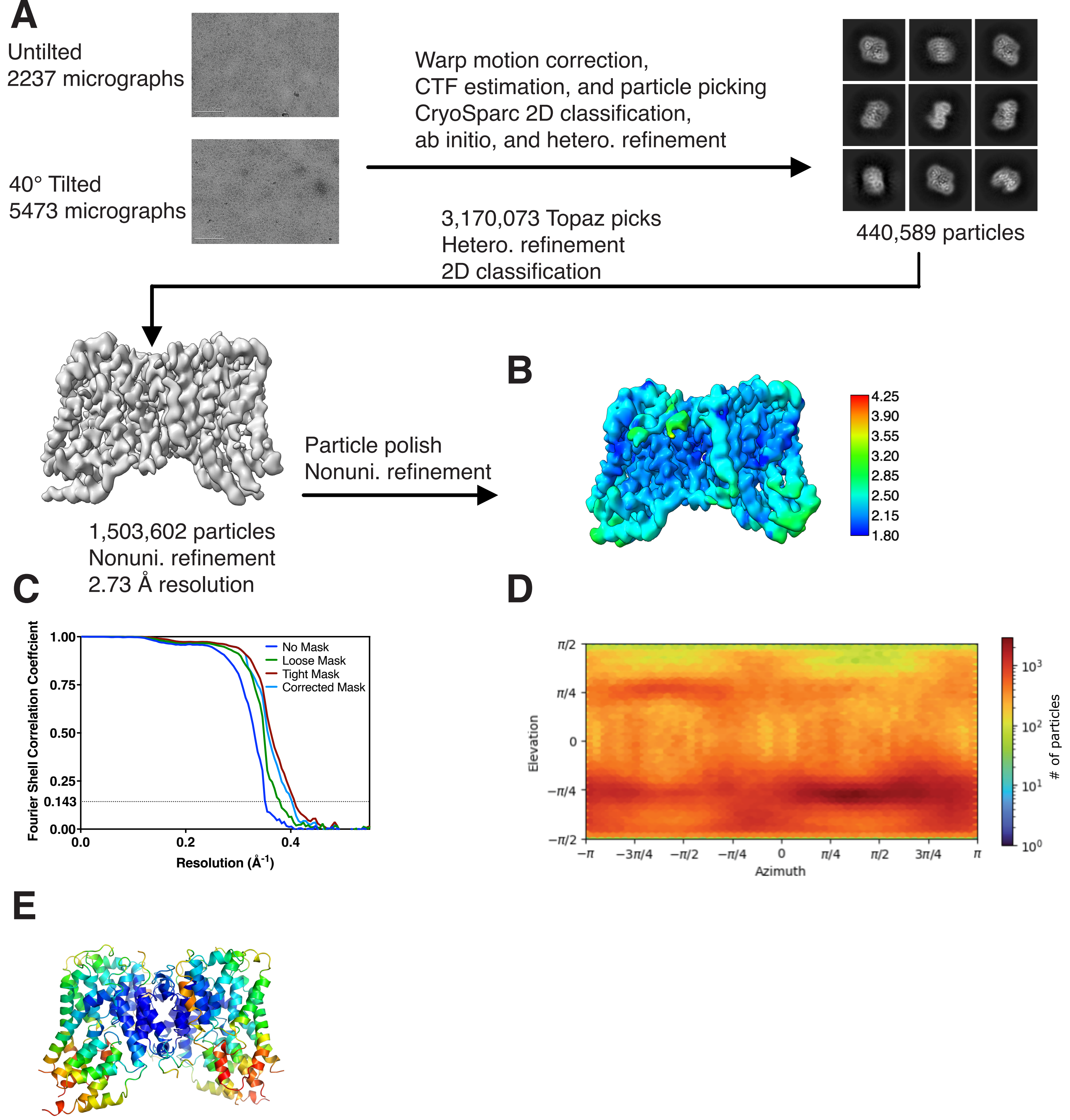


**Supplementary Figure 3. Cryo-EM data processing of the NaCT in Choline Chloride structure.**
A. Cryo-EM data processing workflow for the NaCT in choline chloride dataset. B. Local resolution map of the NaCT in choline chloride map. C. FSC curve for the NaCT in choline chloride map. D. Angular distribution of particles in the NaCT in choline chloride map. E. Model of NaCT in choline chloride, colored by the relative atom B-factors.
